# Identification, Localization and Expression of NHE Isoforms in the Alveolar Epithelial Cells

**DOI:** 10.1101/2020.09.03.280677

**Authors:** Safa Kinaneh, Yara Knany, Emad Khoury, Reem Ismael-Badarneh, Shadi Hammoud, Gidon Berger, Zaid Abassi, Zaher S. Azzam

## Abstract

Na+/H+ exchangers (NHEs), encoded by Solute Carrier 9A (SLC9A) genes in human, are ubiquitous integral membrane ion transporters that mediate the electroneutral exchange of H+ with Na+ or K+. NHEs, found in the kidney and intestine, play a major role in the process of fluid reabsorption together via Na+,K+-ATPase pump and Na+ channels. Nevertheless, the expression pattern of NHE in the lung and its role in alveolar fluid homeostasis has not been addressed. Therefore, we aimed to examine the expression of NHE specific isoforms in alveolar epithelium cells (AECs), and assess their role in congestive heart failure.

Three NHE isoforms were identified in AEC and A549 cell line, at the level of protein and mRNA; NHE1, NHE2 and mainly NHE8, the latter was shown to be localized in the apical membrane of AEC. Treating A549 cells with angiotensin (Ang) II for 1 and 3 hours displayed a significant reduction in NHE8 protein abundance and to lesser extent at 5 hours; however, there was no effect at 24 hours. Moreover, A549 treated overnight with Ang II downregulated NHE8 protein abundance.

CHF rats held for 1 week had increased abundance of NHE8 compared to sham operated rats. However, lower abundance of NHE8 was observed in CHF rats held for 4 weeks.

Herein we show, for the first time, the expression of a novel NHE isoform by AEC, namely NHE8. Besides being negatively affected by Ang II, NHE8 protein levels were distinctly affected in CHF rats, which may be related to CHF severity.

## Introduction

Alveolar fluid clearance has been shown to be an important mechanism in keeping the airspaces free of edema in both cardiogenic and non-cardiogenic states [1,2]. There is a large body of evidence that the removal of alveolar fluid is attained by the alveolar epithelial active sodium transport; by which sodium passively enters the alveolar epithelial cells (AEC) via apical amiloride-sensitive Na^+^ channel (ENaC) or other Na^+^ channels and is pumped out of the cells by basolateral Na^+^, K^+^-ATPase, an energy consuming process. Following sodium transport, water is extruded from the alveolar airspaces [3–5]. It has been shown that the survival of acute lung injury patients, directly correlated with the rate of alveolar fluid clearance [6].

The sodium hydrogen exchanger (NHE) family includes several isoforms, which have different characteristics, including cell-compartment localization, plasma membrane distribution and organ-dependent function [7,8].

The evidence regarding NHE expression and role in the lungs, particularly in alveolar epithelial cells, is scarce. According to the evidence on NHEs in the kidney and intestine; water reabsorption is achieved by the function of Epithelial Na^+^ channel (ENaC), Na^+^,K^+^-ATPase pump along with Na^+^/H^+^ Exchangers (NHEs) [9]; thus, it is conceivable to assume that NHE may contribute to this transport in the lung, specifically AEC. Therefore, the major objective of this work is to address whether NHE isoforms are expressed in alveolar epithelial cells and to evaluate their role in alveolar epithelial active sodium transport in healthy and congestive heart failure (CHF) rats.

## Materials and Methods

### Animals

Experiments were performed on adult male Sprague Dawley rats (Harlan Laboratories Ltd. Jerusalem, 275-350 g). Rats were provided water and food *ad libitum*. The use of animals in the present study was approved by the Technion Institutional Animal Care and Use Committee and it is according to NIH guidelines.

Congestive heart failure rat model was induced by surgically creating Aorto – Caval fistula (ACF) as previously described [10]. The abdomens of Sham controls were opened and sutured back again without creating an A – V fistula. After one or four weeks of the surgery, rats were sacrificed; lungs were collected and stored at -80° C.

### Alveolar epithelial type II cell isolation

Alveolar epithelial type II cells were isolated from the lungs of the various experimental groups based on a method described by Dobbs et al. [11]. Briefly, lungs were instilled and digested with elastase containing solution. Following this, the lungs’ lobes were chopped to release detached cells. Then, cells were plated on IgG coated plates to get rid of contaminating macrophages. The non-adherent cells, mainly AEC, were recovered and plated for further experimentations. The purity of AEC was assessed by modified papanicolaou stain based on the presence of dark blue inclusions. Cell viability was assessed by trypan blue exclusion (>95%).

### Reverse transcriptase PCR (RT-PCR)

RNA purification of AECII and A549 cells was performed using Qiagen RNeasy Mini kit; While RNA purification of lung tissue was achieved using Tri-Reagent method. All steps were done according to the manufacturer’s instructions. The isolated RNA was converted to cDNA using Maxima first strand cDNA kit.

The primers used for RT-PCR are listed in Table 1. cDNA template (10-100ng), forward and reverse primers (0.4µM) and Taq PCR Master mix were added all together to a final volume of 25 µl and placed in thermo cycler machine under the following thermal conditions: 95°C 5 min; 30 cycles of denaturing at 95°C for 30 sec, annealing at 58-62°C for 30 sec, and extension at 72°C for 1min; a final extension at 72°C for 5 min. Following the amplification procedure, PCR products were separated on 1.5% agarose gel with 100 bp DNA ladder and visualized with UV light.

**Table 1.**
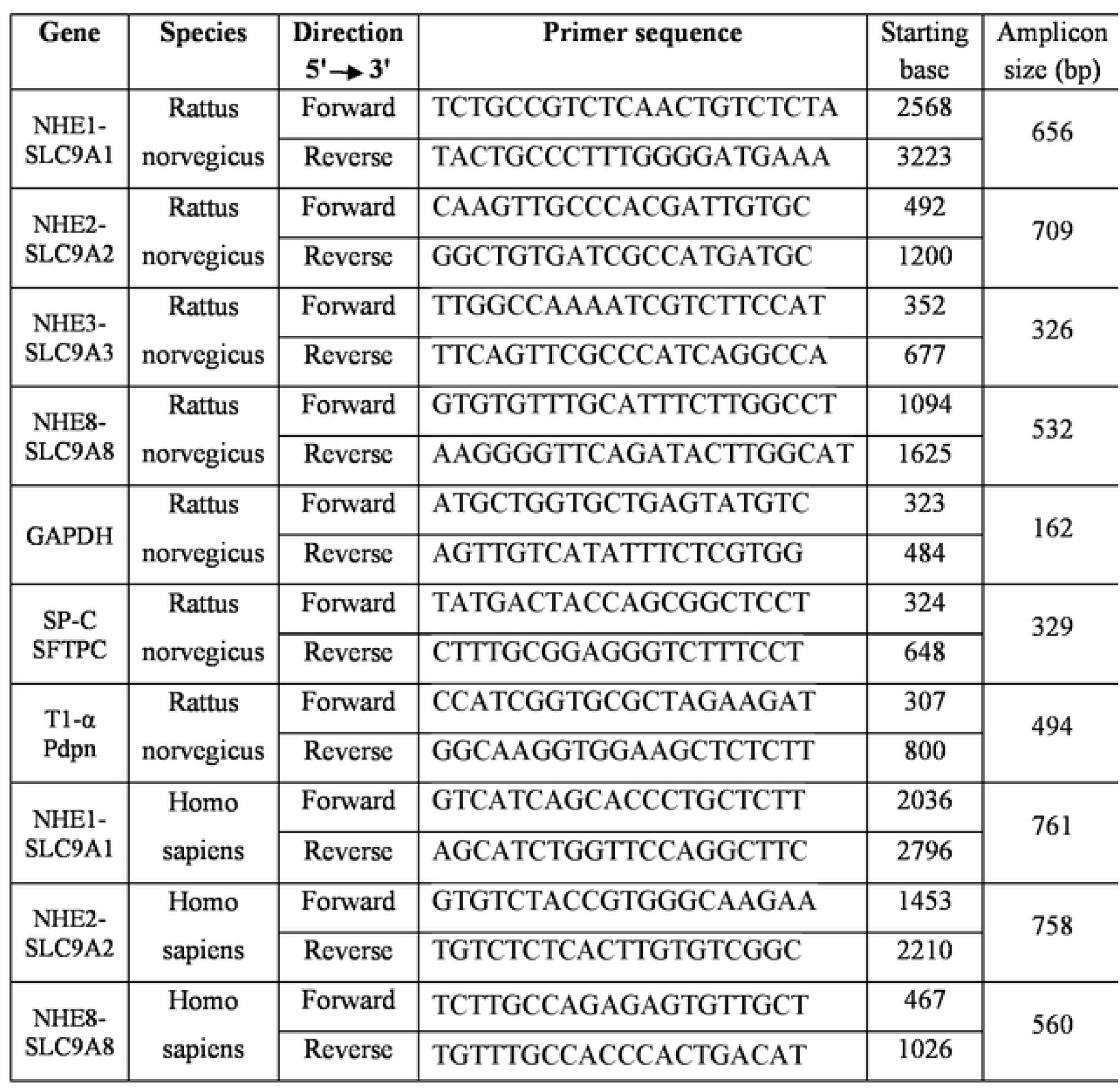

### Cell Lysate and Western Blot Analysis

Equal amounts of protein from lung homogenate, total AEC lysate, or Basolateral plasma membranes (BLMs), were resolved by 10% SDS-PAGE and analyzed by immunoblotting with specific antibodies against, NHE8 isoform and GAPDH, used as internal control.

### Immunofluorescence

A549 or AEC were fixed with 4% then washed with DPBS, incubated with a blocking solution and later incubated overnight at 4° C with primary antibody against NHE8; then, cells were incubated with secondary antibody Alexa-Fluor 594 Donkey anti mouse IgG, washed, and finally mounted with Dapi Immunomount. Sections were visualized using Zeiss Axio observer inverted microscope system.

### Statistical Analysis

Data were presented as mean ± SEM; n is the number of animals in each study group. One way analysis of variance was used when multiple comparisons are made followed by a multiple comparison test (Tukey) when the F statistic indicated significance. To analyze paired data, we used unpaired t-test to assess the differences between the study groups. Results were considered significant when p < 0.05.

The Kolmogorov-Smirnov test was used to analyze the normality of the groups. The Levene’s test was used for comparing the equality of variances. The Student t-test for independent groups was used to compare between two study groups. Two-tailed p value of 0.05 or less was considered to be statistically significant.

## Results

### Identification of Na^+^/H^+^ exchanger isoforms expressed in alveolar epithelial and A549 cells

Our major focus was on NHE isoforms that are primarily localized to cell membranes, thus can potentially contribute to the alveolar active sodium transport, and eventually alveolar fluid clearance. Among these are NHE1-5 and NHE8. NHE5, however, was not included in our experiments as it is reported to be exclusively expressed in the brain. By using RT-PCR and targeted primers to each isoform, the expression of NHE1 and NHE8 was confirmed in isolated AEC (Figure 1A). Moreover, NHE2 was found to be expressed only following cells incubation for 24 hours (Figure 1B); this observation might be attributed to the differentiation of AEC type II into type I. Similarly, these exchangers, namely NHE1, NHE2 and NHE8 are expressed in A549 cell line, known to have characteristic features of AECII (Fig. 1C). Surprisingly, NHE3 expression was not demonstrated in neither cell types, whereas the expression of unexpected novel isoform, namely NHE8, was established.

**Figure 1:**
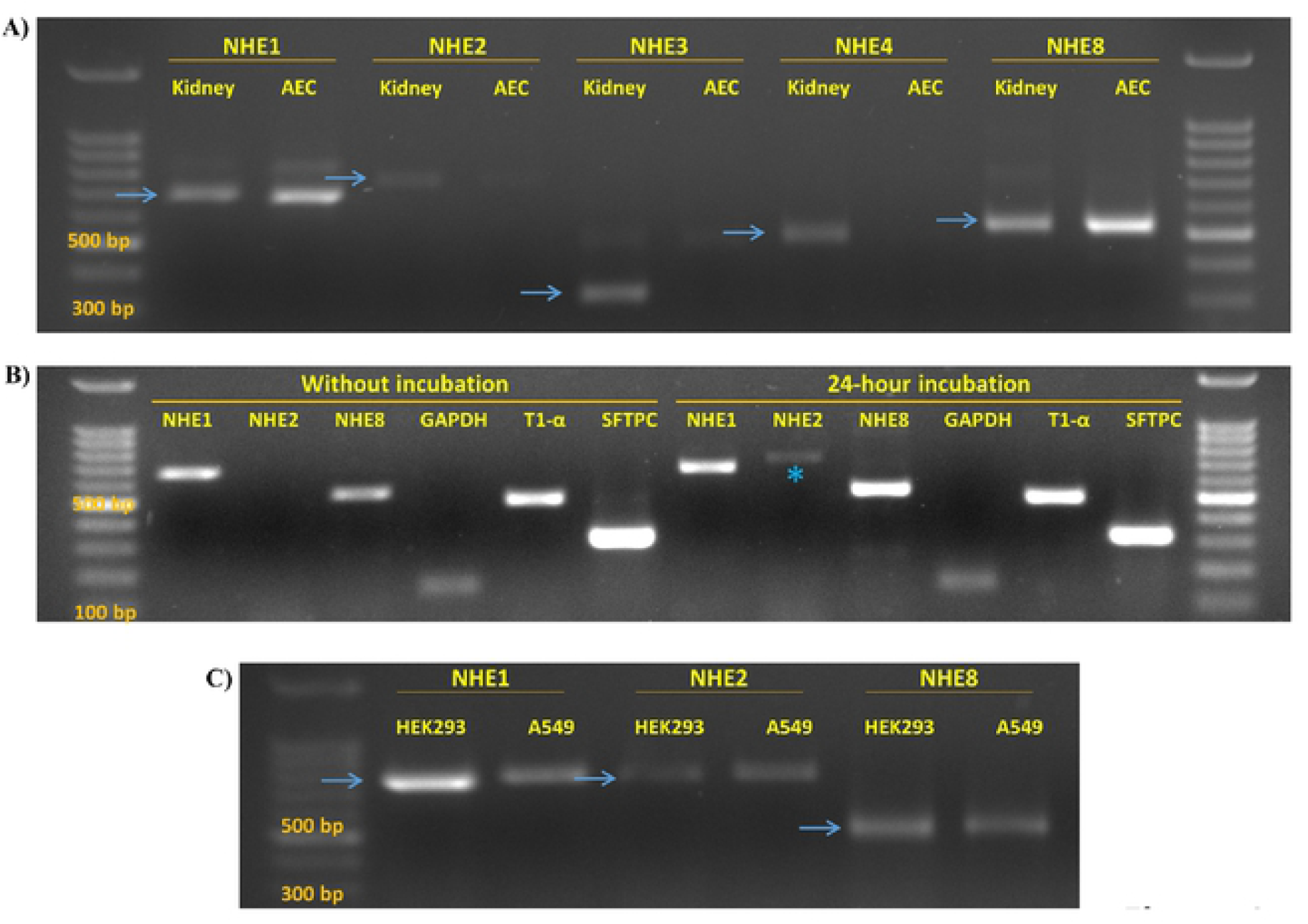
(A) RT-PCR showing the expression of NHE isoforms in isolated AEC as compared to kidney sample. Positive signals of NHE1 and NHE8, but not NHE2 and NHE3, were detected in AEC. (B) RT-PCR that illustrates NHE isoforms expression in AEC that were/or not 24-hour incubated. The asterisk indicate NHE2 expressed in AEC following incubation, but was absent when AEC were freshly assessed. (C) RT-PCR demonstrating the expression of NHE1, NHE2 and NHE8 in A549 and HEK293 that served as positive control. The blue arrows point to the expected product length. AEC - alveolar epithelial cells. NHE – Na^+^/H^+^ Exchanger. GAPDH – Glyceraldehyde 3-phosphate dehydrogenase. T1-α – AECI marker, SFTPC – Surfactant Protein C, AECII marker.

Based on previous reports, NHE8 might be localized to intracellular compartments or to the plasma membrane. Immunofluorescence staining to NHE8 in AEC and A549 cells indicated plasma membrane localization of this exchanger with some cells showing polarity abundance (Figure 2A-C). NHE8 protein was not detected in BLM of lung fractions, whereas Na^+^, K^+^-ATPase is normally localized (Figure 2D).

**Figure 2:**
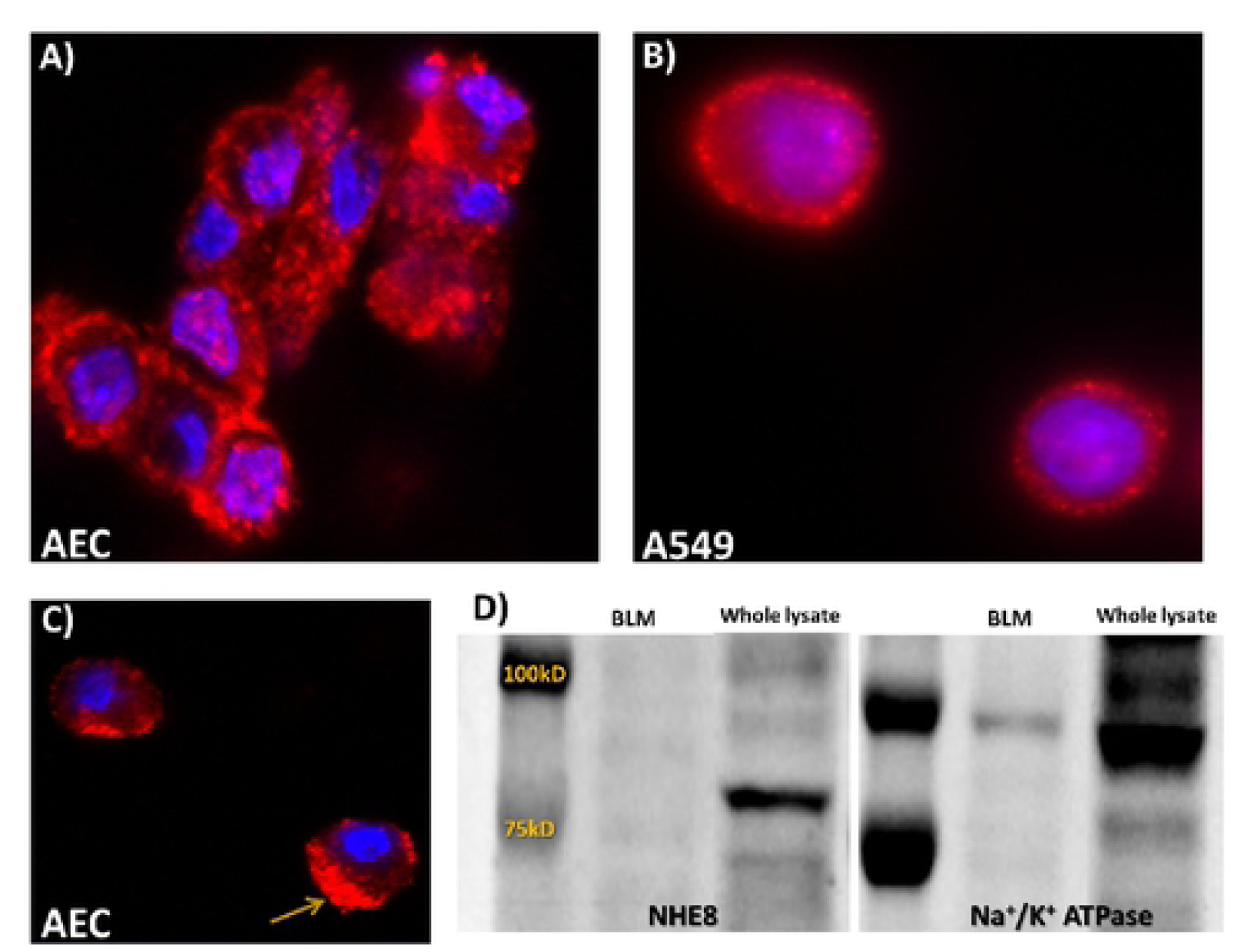
Immunofluorescence staining of NHE8 stained with Cy3 (Red) in isolated AEC (A and C) and A549 cell line (B), showing strong staining at the plasma membrane and a polar distribution in part of the cells (C). Nuclei were stained with Dapi (blue). (D) Western blot to BLM and whole lung lysate showing no existence of NHE8 in BLM fraction; yet, Na^+^/K^+^ ATPase used as a marker of basolateral membranes do exist. BLM – basolateral membranes. NHE – Na+/H+ Exchanger.

### NHE8 levels are decreased in A549 cells following treatment with Angiotensin II

Recently, we have shown that angiotensin II (Ang II) impaired the ability of the lungs to clear edema by downregulating alveolar active Na^+^ transport [12]. Notably, there are no known inhibitors or activators of NHE and assuming that NHE8 may contribute to fluid clearance; we investigated whether it is affected by Ang II. For this purpose, we examined the Ang II effect on NHE8 protein expression in a time - dependent manner. We treated A549 cells with Ang II (10^−8^ M) for 1, 3, 5 and 24 hours. Treating cells for 1 and 3 hours resulted in a significant reduction in NHE8 protein levels, and to a lesser extent following 5 hours treatment, as compared to untreated cells. In contrast, no change in NHE8 protein levels was detected following 24 hours of Ang II treatment (Figures 3A-D, 3F-I). When A549 cells were treated overnight (∼18h) with Ang II at concentrations of 10^−10^ M and 10^−12^ M; NHE8 abundance was downregulated in both concentrations as compared to untreated cells (Figures 3E and J). Notably, this decrease did not reach statistical significance due to the small number of rats in each group (N = 2).

**Figure 3:**
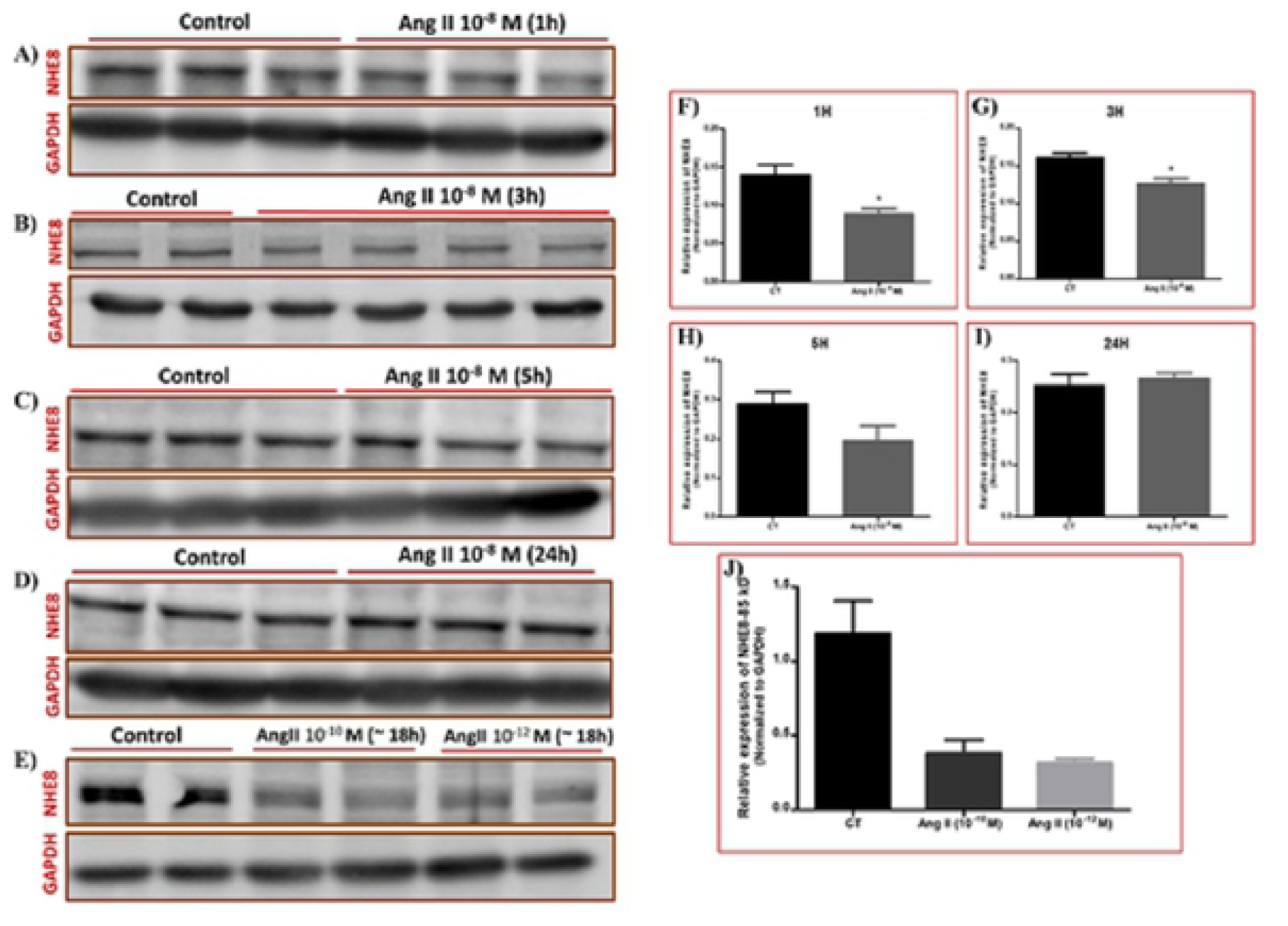
Effect of Angiotensin II treatment on NHE8 protein abundance in A549 cells. A concentration of 10^−8^ M of Ang II were added to A549 cells medium for 1, 3, 5, and 24 hours. NHE8 protein abundance, as assessed by western blot, significantly decreased following Ang II treatment for 1 and 3 hours (A, B, F and G), slightly declined following 5 hours treatment (C and H) and didn’t change after 24 hour treatment (D and J). A549 cells were treated overnight (∼ 18h) with Ang II at two concentrations (10^−10^ M and 10^−12^ M, n=2 in each), which resulted in decreased NHE8 protein levels as shown by western blot (E) and the relevant quantification after normalizing to GAPDH (J). P_Value <0.05. Ang II – Angiotensin II. CT – Control. NHE – Na+/H+ Exchanger. GAPDH – Glyceraldehyde 3-phosphate dehydrogenase. The bars represent mean ± SEM.

### NHE8 protein abundance in lung tissue of CHF-operated rats

Based on the up mentioned finding of NHE8 abundance in AEC, and its plasma membrane occurring profile, we assume that NHE8 has an important role in alveolar fluid clearance; therefore, we were interested to explore whether its expression is modified in a model of congestive heart failure (CHF) as compared to sham rats. We performed western blot to evaluate NHE8 protein expression in lungs, one (CHF-1w) and four (CHF-4w) weeks following ACF procedure, as compared to sham operated rats. CHF-1w rats exhibited an increased NHE8 protein abundance compared to sham-1w rats, yet this elevation was not significant, possibly due to high variability of edema severity among CHF rats (Figures 4 A and C). However, NHE8 immunoreactive levels were significantly decreased in CHF-4w rats as compared to sham 4-w rats (Figures 4 B and D). This distinct expression suggests that NHE8 expression may be related to the severity and progression of CHF.

**Figure 4:**
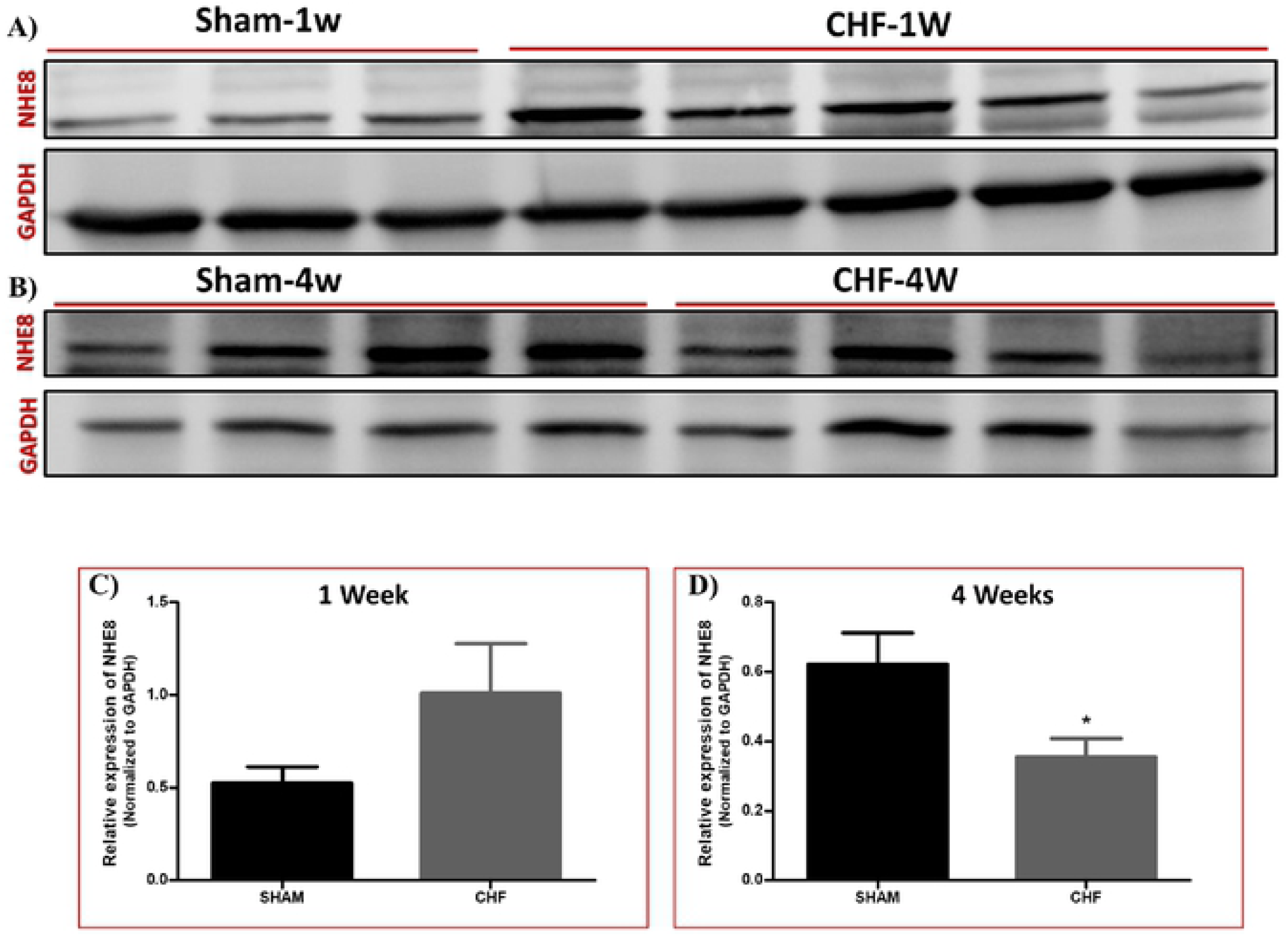
NHE8 immunoreactive levels in the lungs of CHF-1w and CHF-4w rats. (A) Western blot showing increased NHE8 protein levels in CHF-1w (n=5) as compared to their Sham-1w (n=3). (C) Quantification of NHE8; normalized to GAPDH. (B) Western blot showing decreased NHE8 protein levels in CHF-4w (n=4) as compared to their Sham-4w (n=4). (D) Quantification of NHE8; normalized to GAPDH. *P_Value <0.05. The bars represent mean ± SEM. CHF – Congestive Heart Failure. NHE – Na+/H+ Exchanger. GAPDH – Glyceraldehyde 3-Phosphate Dehydrogenase.

## Discussion

Keeping alveolar fluid cavity free of edema is essential for efficient gas exchange. The molecular basis underlying alveolar fluid clearance (AFC) process involves active sodium transport from the apical to the basolateral side of the alveolar epithelium, generating an osmotic gradient that drives water movement and clearance. Mainly, ENaC and Na^+^, K^+^-ATPase are considered to be the main players in this process [3,13,14].

Alveolar fluid clearance process is equivalent to the kidney in terms of water and sodium reabsorption. Yet, in the kidney, another participant, alongside with ENaC and Na^+^, K^+^-ATPase, is involved; that is NHE3. It accounts for most of the sodium reabsorption in the proximal tubules [15]. However, very little is known regarding NHE expression in the lung and particularly in alveolar epithelial cells. Since ENaC contributes to only 40-60% of sodium transport in most species [16], the rest of the sodium entry must be driven by other pathways. Based on this evidence, we were intrigued to investigate the potential role of NHEs in sodium transport, their expression pattern in alveolar epithelial cells, and to evaluate their involvement in pulmonary edema pathology and AFC process. We hypothesized that at least one NHE isoform might be expressed in alveolar epithelial cells that may contribute to vectorial sodium movement.

NHE is a large family containing about nine distinct isoforms. NHE3 has distinctive characteristics and is directly involved in water reabsorption in the kidney, as mentioned previously. Thus, we first, investigated its expression in AEC; however, there was no evidence for NHE3 expression in isolated AEC; an observation that was supported by RT-PCR experiments (Fig. 1A). Therefore, other NHE isoforms known to be localized to the plasma membrane were investigated. Based on previous reports [17,18]; four NHE isoforms were studied; NHE-1, 2, 4, and 8. Our study demonstrated the expression of NHE1 and NHE8 in AEC (Figure 1A), whereas NHE2 and NHE4 expression was not evident.

Notably, while manifesting AECII differentiation properties, the isolated cells were divided into two groups; a freshly processed group and a 24-hour incubation group. NHE isoforms analysis with RT-PCR, demonstrated the expression of NHE2 only in the incubated cells (Figure 1B). Conceivably, this observation may be related to the assumption that NHE2 is localized in AECI.

The AECII cell-like cell line, A549, was used to validate NHE isoforms expression pattern previously described in isolated AEC. Using RT-PCR, NHE1, NHE8 and NHE2 expression was observed in A549 cells as well (Fig. 1C). This set of experiments demonstrated, to our knowledge, for the first time, that NHE8 is expressed in AEC. It is noteworthy that, its expression was confirmed in the kidney and intestine. Obviously, NHE8 expression is remarkably regulated especially in the neonatal intestinal brush border; while in the adults it is replaced with NHE3 [19,20]. Based on these intestinal and renal findings, we decided to focus on NHE8 and further investigate its expression pattern, localization within the cells and involvement in the pathogenesis of pulmonary edema. Therefore, a set of experiments was performed to characterize the localization and regulation of NHE8. Immunofluorescence staining of NHE8 displayed its presence in the plasma membrane of AEC and A549 cells, with indication to polarized abundance (Figures 2A-C). Western blotting to isolated basolateral membranes of AEC was negative for NHE8 expression (Fig. 2D). These findings support the assumption that NHE8 is localized on the apical side of AEC. Unfortunately, there is no known selective inhibitor for NHE8 since it is not much studied and only recently has been discovered. Therefore, we decided to bypass this obstacle and study the response of this isoform to hormones and factors known to impair the ability of the lungs to clear edema.

Angiotensin II, the major component of renin-angiotensin aldosterone system (RAAS), has an important role in the regulation of arterial pressure, salt balance and body fluid volume homeostasis. Furthermore, its levels are increased in heart failure [21]. Recently, our group has shown that Ang II impaired AFC, partly by down-regulating Na^+^, K^+^-ATPase protein levels [12]. Assuming that NHE8 participates in vectorial sodium transport, we hypothesized that Ang II may affect NHE8 protein expression. Therefore, we conducted an experiment in which A549 cells were treated with Ang II (10^−8^ M) and examined NHE8 protein changes over different periods of time-1, 3, 5 and 24 hours. As shown in Fig. 3, Ang II decreased the levels of NHE8 mature protein at 1, 3, and 5 hours; while there was no effect following 24 hours of treatment, which might be explained by Ang II receptor desensitization. Another set of experiments, included overnight treatment of A549 with Ang II at concentrations of 10^−10^ M and 10^−12^ M, showed decreased NHE8 abundance as compared to untreated cells (Figures 3 E and J). Based on these findings, we concluded that Ang II downregulated NHE8 protein abundance in a dose- and time-dependent manner. Notably, since Ang II treatment experiments involved cell incubation for up to 24 hours which might trigger AECI differentiation and in order to minimize the number of sacrificed animals, we favored with the use of A549 cells in this set of experiments, and not isolated AEC.

Furthermore, we investigated NHE8 expression in CHF rats, using ACF rat model. Rats were sacrificed 1 week (CHF-1w) or 4 weeks (CHF-4w) following ACF procedure. NHE8 was distinctly expressed in CHF rat lungs, in which CHF-1w NHE8 levels were increased, while in CHF-4w, NHE8 levels were decreased, as compared to sham-rats (Fig. 4). This distinct expression pattern of NHE8 might suggest a protective role of NHE8 in CHF-1w that is driven by the need to clear excessive lung fluids; while the decreased levels in CHF-4w, might be a result of CHF severe condition, were even protective pathways are badly damaged.

The main limitation of this study was our inability to directly address NHE8 effect on AFC due to the lack of specific inhibitors or NHE8 knock-out mice models. Acute lung injury and deterioration to ARDS is the manifestation of the recently erupted global pandemic, the novel coronavirus SARS-CoV-2 (COVID-19 disease). Therefore, potential involvement of NHEs in general and NHE8 in particular in the pathogenesis of SARS-CoV-2 induced ARDS, worth further evaluation and may contribute to the development of new therapeutic modalities [22,23].

In summary, herein we report, for the first time, the presence of the NHE8 isoform in alveolar epithelial cells. It is presumably localized to the apical membrane of AEC with consequent contribution to the vectorial sodium transport in alveolar epithelial cells. We also showed that Ang II regulates NHE8 protein abundance in a dose- and time-dependent manner, and that NHE8 protein levels are distinctly regulated in CHF-rats (Figure 5).

**Figure 5:**
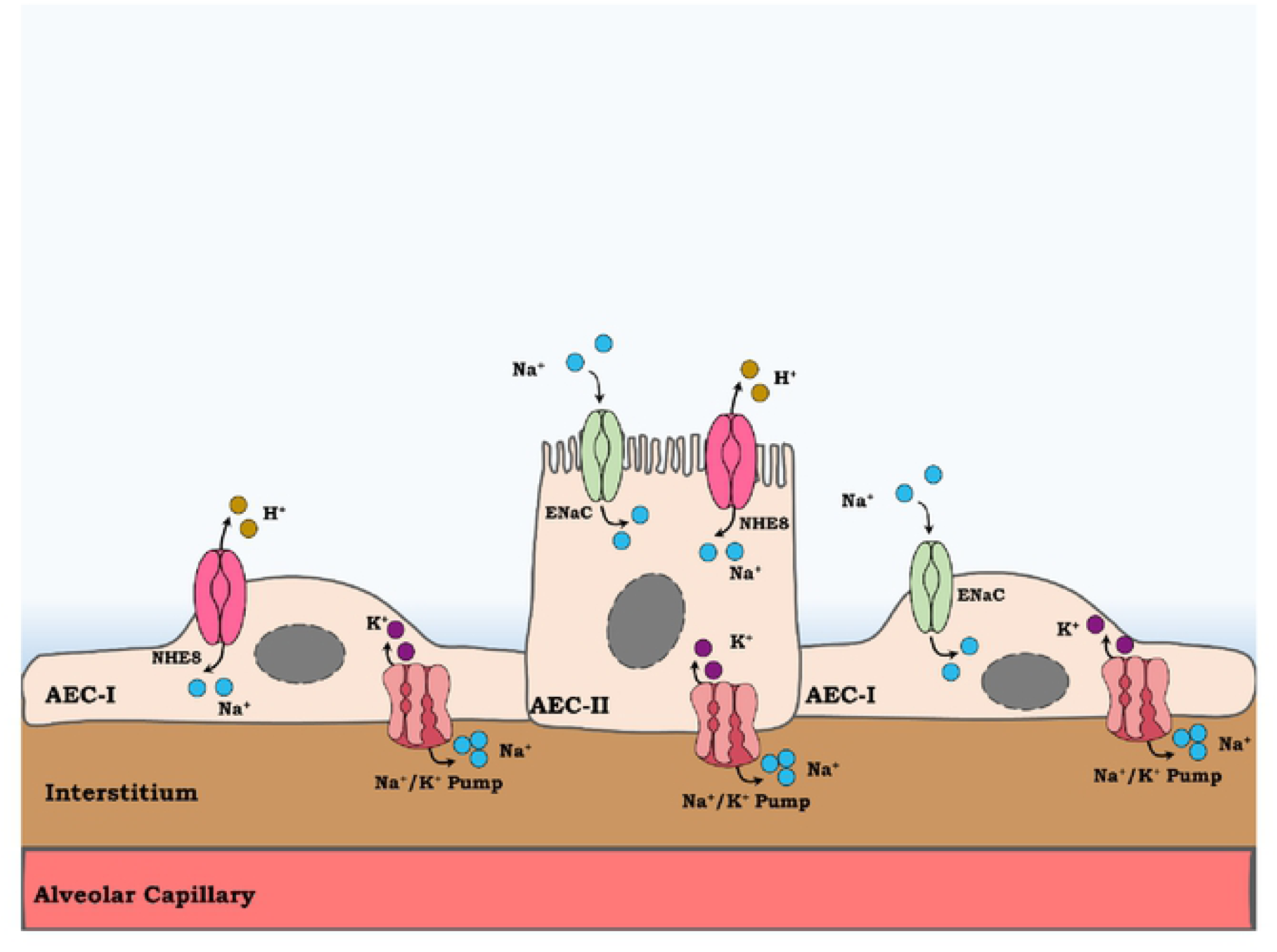
a schematic illustration of NHE8 role in sodium vectorial transport. NHE8 may be located to Alveolar epithelial cells type I (AECI) and II (ACEII), where in the latter it is localized to the apical membrane. By exchanging extracellular Na^+^ with intracellular H^+^, NHE8 contribute to Na^+^ vectorial transport along with epithelial sodium channel (ENaC). The entered sodium then is pumped out of alveolar cells by Na^+^, K^+^ ATpase pump. Sodium transport from alveolar space to the interstitium is accompanied by water movement and so edema clearance.

## Bibliography

1. Matthay MA. Resolution of pulmonary edema thirty years of progress. Am J Respir Crit Care Med 2014;189: 1301–1308. doi: 10.1164/rccm.201403-0535OE

2. Hochberg I, Abassi Z, Azzam ZS. Patterns of alveolar fluid clearance in heart failure. Int J Cardiol 2008;130: 125–130. doi: 10.1016/j.ijcard.2008.03.015

3. Vadasz I, Raviv S, Sznajder JI. Alveolar epithelium and Na,K-ATPase in acute lung injury. Intensive Care Med 2007;33: 1243–1251.

4. Sznajder JI, Factor P, Ingbar DH. Invited review: lung edema clearance: role of Na(+)-K(+)-ATPase. J Appl Physiol 2002;93: 1860–6.

5. Berthiaume Y, Folkesson HG, Matthay MA. Lung Edema Clearance: 20 Years of Progress Invited Review: Alveolar edema fluid clearance in the injured lung. J Appl Physiol 2002;93: 2207–2213. doi: 10.1152/japplphysiol.01201.2001

6. Ware LB, Matthay MA. Alveolar fluid clearance is impaired in the majority of patients with acute lung injury and the acute respiratory distress syndrome. Am J Respir Crit Care Med 2001;163: 1376–1383.

7. Fliegel L, Frohlich O. The Na+/H+ exchanger: An update on structure, regulation and cardiac physiology. Biochem J 1993;296: 273–285. doi: 10.1042/bj2960273

8. Fliegel L. Regulation of myocardial Na+/H+ exchanger activity. Basic Res Cardiol 2001;96: 301–305. doi: 10.1007/s003950170036

9. Slepkov ER, Rainey JK, Sykes BD, Fliegel L. Structural and functional analysis of the Na+/H+ exchanger. Biochem J 2007;401: 623–633. doi: 10.1042/BJ20061062

10. Khoury EE, Kinaneh S, Aronson D, Amir O, Ghanim D, Volinsky N, et al. Natriuretic peptides system in the pulmonary tissue of rats with heart failure: Potential involvement in lung edema and inflammation. Oncotarget 2018;9: 21715–21730. doi: 10.18632/oncotarget.24922

11. Dobbs LG. Isolation and culture of alveolar type II cells. Am J Physiol 1990;258: L134–47.

12. Ismael-Badarneh R, Guetta J, Klorin G, Berger G, Abu-Saleh N, Abassi Z, et al. The role of Angiotensin II and cyclic AMP in alveolar active sodium transport. Bader M, editor. PLoS One 2015;10: 1–13. doi: 10.1371/journal.pone.0134175

13. Matalon S, Hardiman KM, Jain L, Eaton DC, Kotlikoff M, Eu JP, et al. Regulation of ion channel structure and function by reactive oxygen-nitrogen species. Am J Physiol Lung Cell Mol Physiol 2003;285: L1184–9.

14. Matthay MA, Folkesson HG, Clerici C. Lung epithelial fluid transport and the resolution of pulmonary edema. Physiol Rev 2002;82: 569–600.

15. Vallon V, Schwark JR, Richter K, Hropot M. Role of Na+/H+ exchanger NHE3 in nephron function: Micropuncture studies with S3226, an inhibitor of NHE3. Am J Physiol - Ren Physiol 2000;278: F375–F379. doi: 10.1152/ajprenal.2000.278.3.f375

16. Mutlu GM, Sznajder JI. Mechanisms of pulmonary edema clearance. Am J Physiol Lung Cell Mol Physiol 2005;289: L685–95.

17. Brett CL, Donowitz M, Rao R. Evolutionary origins of eukaryotic sodium/proton exchangers. Am J Physiol - Cell Physiol 2005;288: C223–C239. doi: 10.1152/ajpcell.00360.2004

18. Donowitz M, Ming Tse C, Fuster D. SLC9/NHE gene family, a plasma membrane and organellar family of Na +/H+ exchangers. Mol Aspects Med 2013;34: 236–251. doi: 10.1016/j.mam.2012.05.001

19. Gurney MA, Laubitz D, Ghishan FK, Kiela PR. Pathophysiology of Intestinal Na+/H+ Exchange. CMGH 2017;3: 27–4. doi: 10.1016/j.jcmgh.2016.09.010

20. Becker AM, Zhang J, Goyal S, Dwarakanath V, Aronson PS, Moe OW, et al. Ontogeny of NHE8 in the rat proximal tubule. Am J Physiol - Ren Physiol 2007;293: F255–F261. doi: 10.1152/ajprenal.00400.2006

21. Packer M. The neurohormonal hypothesis: a theory to explain the mechanism of disease progression in heart failure. J Am Coll Cardiol 1992;20: 248–254.

22. Wu F, Zhao S, Yu B, Chen YM, Wang W, Song ZG, et al. A new coronavirus associated with human respiratory disease in China. Nature 2020;579: 265–269. doi: 10.1038/s41586-020-2008-3

23. Singer BD, Jain M, Budinger GRS, Wunderink RG. A Call for Rational Intensive Care in the Era of COVID-19. Am J Respir Cell Mol Biol 2020;63: 132–133. doi: 10.1165/rcmb.2020-0151LE

